# ICAN: interpretable cross-attention network for identifying drug and target protein interactions

**DOI:** 10.1101/2022.08.04.502877

**Authors:** Hiroyuki Kurata, Sho Tsukiyama

## Abstract

Drug–target protein interaction (DTI) identification is fundamental for drug discovery and drug repositioning, because therapeutic drugs act on disease-causing proteins. However, the DTI identification process often requires expensive and time-consuming tasks, including biological experiments involving large numbers of candidate compounds. Thus, a variety of computation approaches have been developed. Of the many approaches available, chemo-genomics feature-based methods have attracted considerable attention. These methods compute the feature descriptors of drugs and proteins as the input data to train machine and deep learning models to enable accurate prediction of unknown DTIs. In addition, attention-based learning methods have been proposed to identify and interpret DTI mechanisms. However, improvements are needed for enhancing prediction performance and DTI mechanism elucidation. To address these problems, we developed an attention-based method designated the interpretable cross-attention network (ICAN), which predicts DTIs using the Simplified Molecular Input Line Entry System of drugs and amino acid sequences of target proteins. We optimized the attention mechanism architecture by exploring the cross-attention or self-attention, attention layer depth, and selection of the context matrixes from the attention mechanism. We found that a plain attention mechanism that decodes drug-related protein context features without any protein-related drug context features effectively achieved high performance. The ICAN outperformed state-of-the-art methods in several respects and revealed with statistical significance that some weighted sites in the cross-attention weight represent experimental binding sites, thus demonstrating the high interpretability of the results.

**Key points:** We created the interpretable cross-attention network (ICAN), which is composed of nn.Embedding of FCS label-encoding vectors of SMILES of drugs and AA sequences of target proteins, cross-attention mechanisms, and a CNN output layer.

ICAN decoded drug-related protein context features without any protein-related drug context features, achieving high prediction performance despite the plain attention mechanism.

In comparison with seven state-of-the-art methods, ICAN provided the highest PRAUC for the imbalanced datasets (DAVIS and BindingDB).

Statistical analysis of attention-weight matrixes revealed that some weighted attention sites corresponded to experimental binding sites, demonstrating the high interpretability achievable with ICAN.

## Introduction

Drug–target protein interaction (DTI) identification is fundamental for drug discovery and drug repositioning, because therapeutic drugs act on disease-causing proteins. However, the DTI identification process often requires expensive and time-consuming tasks, including biological experiments involving large numbers of candidate compounds [1]. To address these problems, intensive efforts have focused on virtual screening or computational prediction of DTIs based on large biological datasets and information available in public databases. A variety of computational approaches have been proposed for DTI prediction [2]. Conventional structure-based approaches that include docking simulations have been studied for decades, but they are limited to proteins for which the three-dimensional structure is known or can be precisely reproduced by molecular dynamics simulations. Similarity-based approaches such as kernel regression [3-5] and matrix factorization [6-8] utilize known drug-target similarity information to infer new DTIs. However, these methods are not applicable to target proteins of different classes for which similarity information is lacking [9]. Feature-based methods used in chemo-genomics approaches compute the descriptors of drugs and proteins as the input data to train machine and deep learning models so that they can accurately predict unknown DTIs as classification models and the associated binding affinity as regression models, respectively. Drug compounds can be represented as types of fingerprints, including extended connected fingerprints (ECFPs) [10] and PubChem [11], one-dimensional (1D) sequences, and molecular graphs (traditionally called two-dimensional [2D] structures). Protein sequences are described as 1D sequences, which are converted into feature vectors using various descriptors, including composition transition distribution [12] and protein sequence composition (PSC) descriptors [13]. In general, machine learning methods, such as random forest and gradient boosting, can be combined with such descriptors to predict DTIs [14-18], where appropriate design and selection of descriptors are critically important.

Recent studies have proposed various end-to-end, deep learning frameworks that integrate sequence representation and model training in a unified architecture. Deep learning–based methods such as deep neural network [19], long short-term memory [20, 21], convolutional neural networks (CNNs) [22], and Transformer [13] have improved the prediction performance of machine learning methods. DeepDTI [9] proposed a deep belief network [23] that uses ECFP2, ECFP4, and ECFP6 [10] to encode drugs and PSC descriptors to encode amino acids (AAs). DeepDTA [22] uses two CNNs that extract the features of the Simplified Molecular Input Line Entry System (SMILES) of drugs and AA sequences from their label-encoding vectors. DeepConv-DTI [24] involves an architecture similar to DeepDTA, which uses CNNs with max pooling layers and fingerprint ECFP4 to encode SMILES. DeepAffinity [25] employs recurrent neural networks to learn the feature vectors of drugs and proteins. From another aspect in which a drug compound is regarded as a molecular graph, GNN-CPI [26] and GraphDTA [27] have implemented graph neural networks (GNNs) [28, 29] and graph CNNs [30, 31], respectively. These predictors separately learn the molecular representations of drugs and proteins and then concatenate their features to calculate a final output, while neglecting the interactions of sub-sequences between drugs and proteins.

Such interactions are widely considered by exploiting the attention mechanisms of Transformer, an encoder-decoder model consisting of multi-headed attention layers and pairwise feed-forward to extract sequence-to-sequence features. TransformerCPI [13] implemented a Transformer that applies drugs and proteins via a sequence-to-sequence or cross-attention mechanism. MolTrans [32] employs two self-attention mechanisms to create an interaction map that integrates the separately generated context features from drugs and proteins. These attention-based methods have suggested a few examples that roughly explain how a high value of attention weight matrixes correspond to experimental binding sites, but they do not prove this in an objective and statistical manner.

To overcome these problems, we developed a new attention-based method that predicts DTIs based on SMILES of drugs and AA sequences of proteins; this method was designated the interpretable cross-attention network (ICAN). We optimized the attention mechanism architectures in terms of cross-attention or self-attention, attention layer depth, and selection of the context matrixes from the attention networks. ICAN outperformed state-of-the-art methods in several respects and revealed with statistical significance that some weighted attention sites represent experimental binding sites, thus demonstrating the distinct interpretability of the method.

## Materials and methods

### Computation framework

The proposed cross-attention model consists of three networks: an embedding layer, an attention layer, and an output layer, as shown in **Fig. 1**. In the embedding layer, the SMILES of drugs and AA sequences of target proteins are separately encoded as the embedding matrixes. These matrixes are input into the cross-attention network to generate the drug-related context matrix of a protein and the protein-related context matrix of a drug. The resulting context matrixes are applied to the output layer of the CNN, which functions as a decoder to capture local feature patterns at different levels, followed by computing of a DTI probability score. The cross-attention considers sub-sequence interactions between a drug and a protein to produce the context matrixes; the CNN uses different filters to extract local sub-sequence patters within the context matrixes. Details in encoding methods and attention-based models are described in **Tables 2** and **3**.

**Figure 1.**
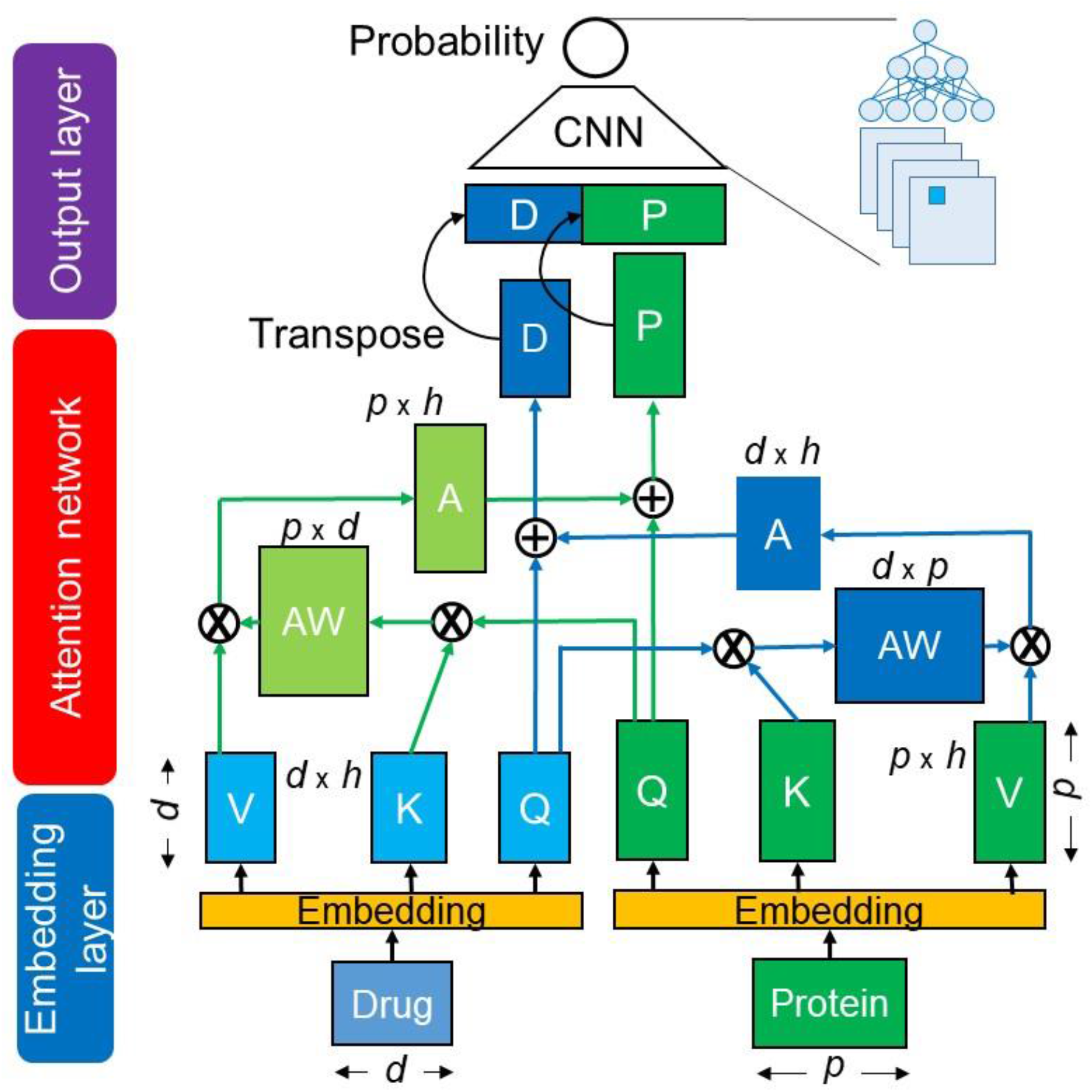
Architecture of the cross-attention-based neural network. It is the CA_DP network (Table 3). Q, K, and V indicate Query, Key, and Value matrixes. AW: attention-weight matrix, A: attention matrix, D: drug-context matrix, P: protein-context matrix, d: length of drug sequence, p: length of protein sequence, h: hidden dimension size.

### DTI dataset

The three widely used datasets were prepared: DAVIS, BindingDB, and BIOSNAP, as shown in **Table 1**. These were the same datasets employed in a previous study [32] and facilitated direct comparison of our proposed method with existing methods. The DAVIS dataset consists of biochemically determined experimental Kd (dissociation constant) values for 68 drugs and 379 proteins [33]. The BindingDB database [34] includes Kd values for 10,665 drugs and 1,413 proteins. DTI pairs with a Kd <30 μM are regarded as positive samples. The MINER DTI dataset from the BIOSNAP collection [35] includes 13,741 DTI pairs for 4,510 drugs and 2,181 proteins from DrugBank [36]. As the BIOSNAP dataset contains only positive DTI pairs, negative DTI pairs are collected and removed from among the unseen pairs [32]. We removed samples for which SMILES was not applicable to RDKit [37].

**Table 1.**
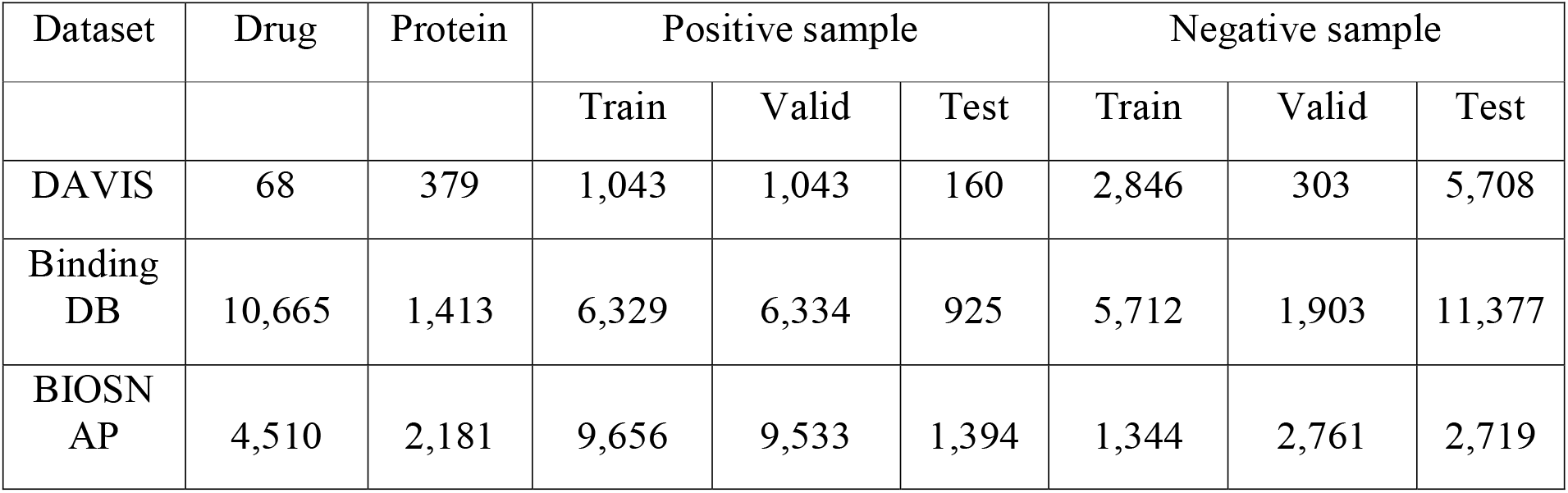
Dataset statistics

Positive DTI pairs were divided into training, validation, and testing sets at a ratio of 7:1:2. For well-shaped training, the number of negative DTI samples was set to the same number as positive samples in the training datasets. The remaining negative pairs were placed in the validation and test datasets. The positive and negative pairs are balanced in the BIOSNAP test dataset; they are imbalanced in the DAVIS and BindingDB test datasets, where the number of negative samples is more than 20-fold and 7-fold greater than that of positive samples, respectively.

### Drug and protein representations

Drugs are represented by the SMILES [38], which is a character sequence of atoms and bonds (e.g., C, N, O, =), generated by depth-first traversal for a molecule graph. The SMILES has the disadvantage that its respective characters do not always correspond to chemically valid molecules. To overcome this problem, SELF-referencIng Embedded Strings (SELFIES) [39] are introduced so that all of the tokens or words of the SELFIES correspond to chemically valid molecules, increasing the reliability of interpretation and rationality of the learning algorithms or mechanisms. It should be noted that a token is the minimum meaningful character set. A target protein is represented as a letter sequence of the 20 standard AAs.

### Frequent consecutive sub-sequence mining

The frequent consecutive sub-sequence (FCS) mining method [32] is used to extract sub-sequences with variable length for both drug compounds and proteins. The FCS algorithm finds recurring sub-sequences across drug and protein sequence datasets. According to the natural language processing–based identification method of sub-word tokens [40, 41], FCS produces a hierarchical set of frequently occurring sub-sequences. After collecting a sequence set of SMILES and AA sequences, the FCS method decomposes or tokenizes the sequences of drugs and proteins into sub-sequences and individual atoms and AAs that are designated a token set. FCS scans the token set to identify the most-frequent consecutive tokens (e.g., X, Y) and updates them with their combined token (XY). FCS repeats the process of scanning, identification, and updating until no frequent token is above a specific threshold or the number of tokens reaches a specific threshold. Frequent sub-sequences are bound into one token, and infrequent sub-sequences are decomposed into short tokens.

### Encoding methods

We paired each token of SMILES FCS, AA FCS, SMILES, SELFIES, and AAs with its corresponding label integer to generate a label-encoding vector. The resulting label-encoding vectors were embedded into token-embedding matrixes by two encoding methods, nn.Embedding (PyTorch) and one-hot encoding. **Table 2** shows the five encoding methods employed: (1) nn.Embedding of FCS embeds the label-encoding vectors of SMILES FCSs and AA FCSs; (2) nn.Embedding of SMILES embeds the label-encoding vectors of SMILES characters and AA letters; (3) nn.Embedding of SELFIES embeds the label-encoding vectors of SELFIES words and AA letters; (4) one-hot encoding of SMILES embeds the label-encoding vectors of SMILES characters and AA letters; and (5) one-hot encoding of SELFIES embeds the label-encoding vectors of SELFIES words and AA letters. Here, we demonstrate how to make a label encoding vector with SMILES characters. SMILES [CN=C=O] is decomposed into a character list: [C, N, =, C, =, O], which is then converted into a label-encoding vector [1, 3, 60, 1, 60, 4, 60], where (C:1, N:3, =:60, O:4).

**Table 2.** Drug and protein encoding methods

| Name | Drug token | Protein token |
| --- | --- | --- |
| nn.Embedding of FCS | SMILES FCS | AA FCS |
| nn.Embedding of SMILES | SMILES character | AA letter |
| nn.Embedding of SELFIES | SELFIES word | AA letter |
| one-hot encoding of SMILES | SMILES character | AA letter |
| one-hot encoding of SELFIES | SELFIES word | AA letter |

“Character” and “word” indicate “a letter or a symbol” and “minimum chemically valid characters”, respectively.

As drug and protein sequences have varying lengths, the maximum sizes of the label-encoding vectors for drugs and proteins were set to 50 and 545, respectively. nn.Embedding embeds the label-encoding vectors of a drug and protein into 50 × 384 and 545 × 384 token-embedding matrixes, respectively. The one-hot encoding method encodes a drug and a protein into 50× (length of the SMILES character list or SELFIES word list) and 545× (length of AA letter list) token-embedding matrixes, respectively.

We also added a positional embedding method to the token-embedding, because the position of words generally plays an essential role in any language grammar, defining the semantics of a sentence. As the token-embedding matrixes do not have any information regarding position of the sub-sequences or tokens, we added position information for the sub-sequences to their features. The position label–encoding vector is embedded by nn.Embedding into the position-embedding matrix, which is added to the token-embedding matrix. The resulting matrix was designated the embedding matrix.

### Cross-attention network

We built multi-head cross-attention networks with an attention head number of *h*. The query of a target protein, 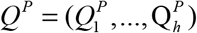, is provided with the embedding matrix of a protein *Y* ^*P*^ by:

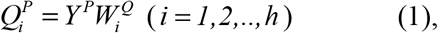

Where 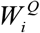 represents the weight matrix. The Key and Value of a drug, 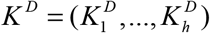 and 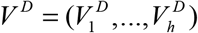, are provided with the embedding matrix of a drug *Y* ^*D*^ by:

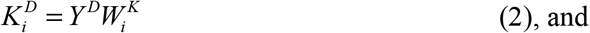

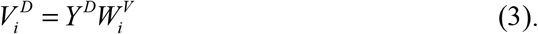

We calculated the dot product of 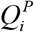 and 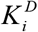 to obtain the attention weight matrix given by:

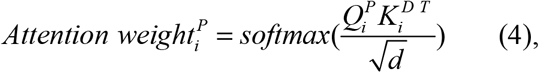

where *d* represents the column dimension of 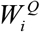 and 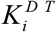 represents the transposed matrix of 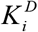 The attention weight matrix is multiplied by 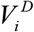 to compute the attention, or head, which is given by:

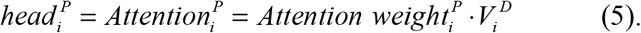

The multi-head attention is expressed by:

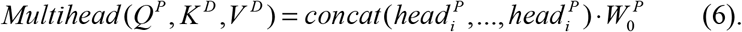

Subsequently, context matrix of protein *C*^*P*^ is given by

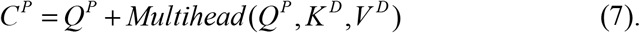

Next to the context matrix of a protein, we computed the following context matrix of a drug, and vice versa.

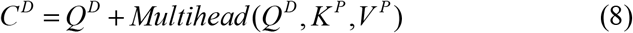

The final context matrixes from the cross-attention networks have three types: consecutively concatenated context matrixes of a drug and protein (*C*^*D*^, *C*^*P*^), context matrix of a drug alone *C*^*D*^, and context matrix of a protein alone *C*^*P*^.

The resulting context matrix was sent to the output layer to obtain a final probability or prediction score. The output layer consists of two sets of 1D-CNN, ReLU function, max pooling layer, and dropout layer, and one linear transformation layer with the sigmoid function. As an alternative output layer, we prepared fully connected layers composed of two sets of a linear transformation layer, ReLU function, and dropout layer, and one linear transformation network with the sigmoid function.

### Evaluation

Six statistical metrics were used to evaluate the prediction performance of the proposed models: sensitivity (recall) (SN), specificity (SP), precision (PR), F1, area under the receiver operating characteristic curve (ROCAUC), and area under the precision and recall curve (PRAUC). SN, SP, PR, and F1 are given by:

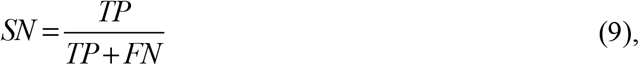

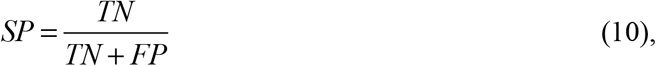

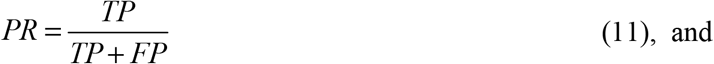

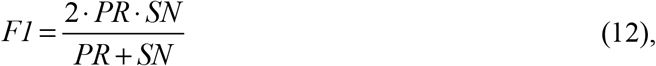

where *TP, TN, FP*, and *FN* denote the numbers of true positives, true negatives, false positives, and false negatives, respectively. We conducted five independent runs to train the models and evaluate them with the test datasets. The statistical metrics were averaged over 5 models. PRAUC was an effective metric on the imbalanced test datasets of DAVIS and BindingDB, where the number of negative pairs was much larger than that of positive pairs.

### Attention site analysis

To demonstrate the interpretability of the attention mechanisms, we investigated the relationship between a high value in the attention weight matrix and the experimental drug-binding sites of target proteins or whether the attention sites correspond to the experimental DTI binding sites. First, we added experimental data regarding drug-binding sites, as derived from sc-PDB [42], to the BIOSNAP test dataset. Consequently, we updated 160 DTIs with their binding sites. Notably, DTI samples with experimental drug-binding sites are limited. Second, we fed the 160 DTIs into the CA_P model (**Table 3**) to generate attention weight matrixes, which were expanded by the protein and drug sequence axes. For each DTI sample, we sorted the attention sites according to value and then selected the top 30 attention sites. Third, we counted the attention sites that corresponded to the experimental binding sites for each target protein, and this value was designated the experimental consistency number. Fourth, to statistically validate the experimental consistency numbers, we artificially generated random binding sites for each target protein, while keeping the number of random sites the same as the number of experimental sites. We then calculated how many times the top 30 attention sites corresponded to the randomly generated binding sites, which was designated the simulated consistency number. This simulation process was iterated 10,000 times for the 160 DTIs to obtain a profile of simulated consistency numbers. Finally, based on the simulated profile, we calculated the z-score of the experimental consistency number between the top 30 attention sites and the experimental binding sites.

**Table 3.** Architectures of different attention-based models

| Method name | Attention mechanism | Attention layer depth | Context matrix | Output layer |
| --- | --- | --- | --- | --- |
| CA_DP | Cross-attention | 1 | Drug+Protein | CNN |
| CA2_DP | Cross-attention | 2 | Drug+Protein | CNN |
| CA3_DP | Cross-attention | 3 | Drug+Protein | CNN |
| CA_DP_FCL | Cross-attention | 1 | Drug+Protein | FCL |
| SA_DP | Self-attention | 1 | Drug+Protein | CNN |
| CA_D | Cross-attention | 1 | Drug | CNN |
| CA_P (ICAN) | Cross-attention | 1 | Protein | CNN |

All of the models employ the nn.Embedding of FCS (Table 2).

### Implementation

All the programs were written in Python. Programming regarding SMILES and SELFIES was written using RDKit [37]. The deep learning programs, including the attention mechanisms and CNN, were implemented in PyTorch [43].

## Results and discussion

### Optimization of neural network structures

There are a number of network architectures in the attention-based methods (**Table 3**). We designed or optimized the network structures of the attention-based network and output layer using the DAVIS dataset with nn.Embedding of the FCS. This optimality of this encoding will be demonstrated later. The attention mechanism selects either of two types: self-attention or cross-attention. The depth of attention layers varies as 1, 2, or 3. The context matrixes resulting from the attention network have three types of concatenated context matrixes: drug and protein, drug context alone, and protein context alone. The output layer employs either of two neural networks: CNN or fully connected layer.

First, we defined CA_DP (**Table 3, Fig. 1**) as the base model that consists of nn.Embedding of the FCS, one-layer cross attention mechanisms for the concatenated matrixes of drugs and proteins, and CNN output layer, and tuned its hyper-parameters. Consequently, we determined the number of multi-head attentions as 4, the learning rate as 0.001, maximum epochs as 50, and training batch size as 128. The Adam optimizer was used. All hyper-parameters are provided in our program. To select the neural network suitable for the output layer, we compared CA_DP with CA_DP_FCL that used the fully connected layer as the output layer, as shown in **Fig. 2** and **Table S1**. CNN presented higher SN, ROCAUC, and PRAUC values than FCL; thus, CNN was selected. **Table S1** indicates the values of all the six metrics including PR and F1. We considered that CNN captured local context patterns with different filters. To investigate the effect of attention layer depth on performance, we compared one layer (CA_DP), two layers (CA2_DP), and three layers (CA3_DP). An increase in the layer number decreased the ROCAUC and PRAUC values, suggesting that complex attention mechanisms do not improve performance. Thus, we selected a one-layer attention mechanism. We compared the cross-attention (CA_DP) and self-attention (SA_DP) mechanisms (**Fig. S1**). The cross-attention mechanism exhibited better performance for the four metrics than the self-attention mechanism. This suggests that interactive cross-attentions between drug and protein features are critically important for prediction. This was not consistent with the observation that the two self-attention mechanisms employed by MolTrans exhibit high prediction performance. We speculate that MolTrans complements the lack of cross-attention between drugs and proteins with an interaction map that integrates drug and protein context features from the self-attention mechanism.

**Figure 2.**
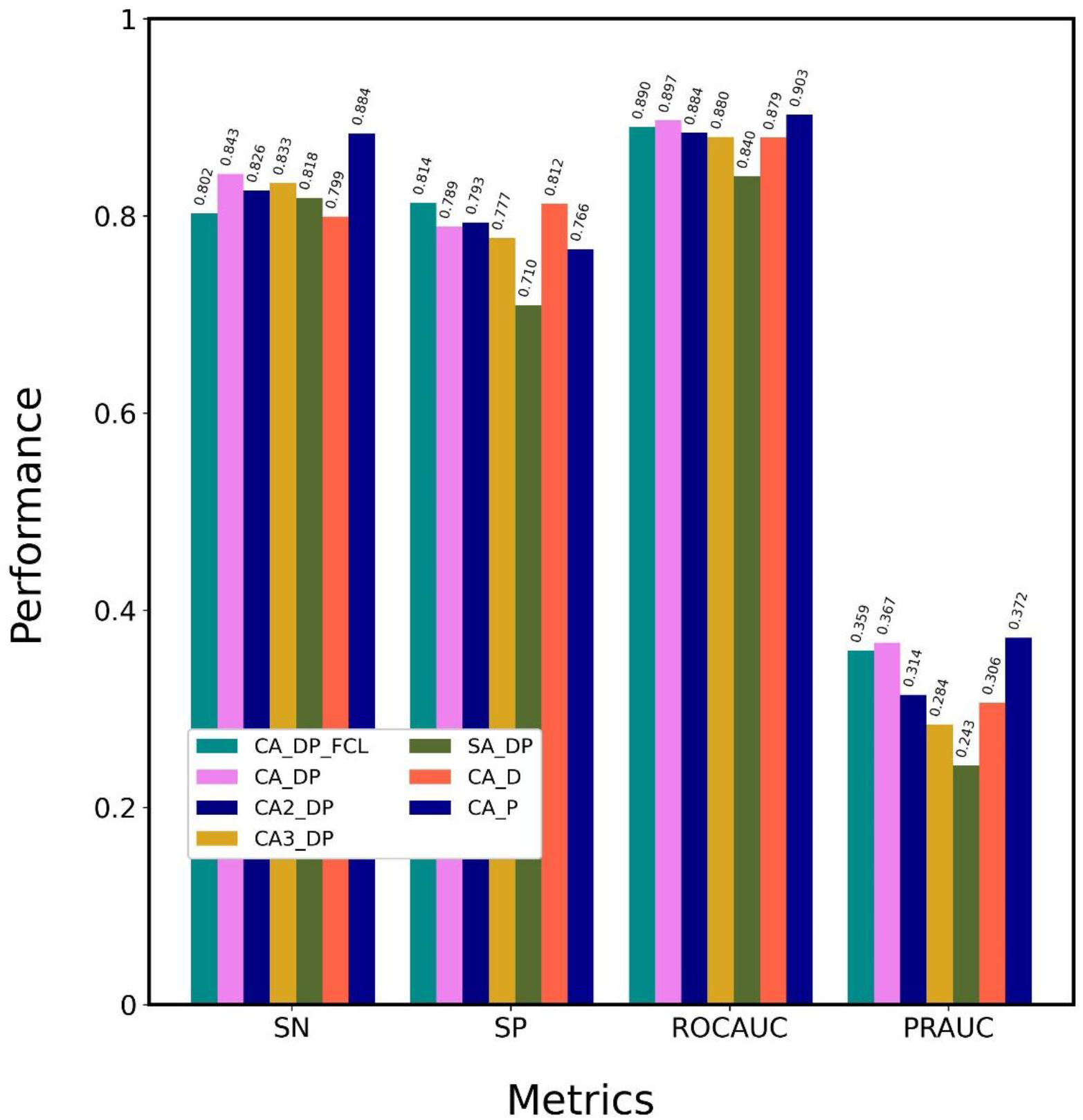
Optimization of the attention-based network architecture. Attention-based models with nn.Embedding of FCS were trained using the DAVIS training dataset and evaluated using the DAVIS test dataset. Details of the methods are shown in Table 3. CA_P was designated ICAN.

To select the optimal context matrixes resulting from the attention mechanisms, we compared CA_DP, CA_D, and CA_P (**Fig. S2**). CA_P provided higher SN, ROCAUC, and PRAUC values than CA_DP and CA_D, suggesting that CA_P decodes drug-related protein context features without any protein-related drug context features. This is similar to TransformerCPI, which decodes protein features based on encoded drug features, whereas CA_P uses a plain one-attention-layer architecture without any pairwise feed-forward. We conclude that the plain attention mechanism, CA_P, is sufficiently effective to achieve high prediction performance.

### Comparison of encoding methods

To demonstrate the superiority of the nn.Embedding of FCS, five CA_Ps with different encoding methods (**Table 2**) were tested on the DAVIS dataset, as shown in **Fig. 3** and **Table S2**: nn.Embeddings of FCS, SMILES, SELFIES, and one-hot encodings of SMILES and SELFIES. nn.Embedding of FCS presented the highest SN, ROCAUC, and PRAUC values. nn.Embeddings of SMILES and SELFIES outperformed the one-hot encodings of SMILES and SELFIES, suggesting that nn.Embedding facilitates attention-based learning to a greater degree than one-hot encoding. This difference is due to the perceptron architecture implemented by nn.Embedding. nn.Embedding of SELFIES provided higher SP, ROCAUC, and PRAUC values than nn.Embedding of SMILES; one-hot encoding of SELFIES provided much higher SN, ROCAUC, and PRAUC values than one-hot encoding of SMILES. This result indicates that SELFIES is more effective than SMILES at learning features, because SELFIES decomposes chemical formulas into chemically meaningful character lists, whereas SMILES does not. These optimization processes confirmed that CA_P with nn.Embedding of FCS is the optimal method, which was then designated ICAN.

**Figure 3.**
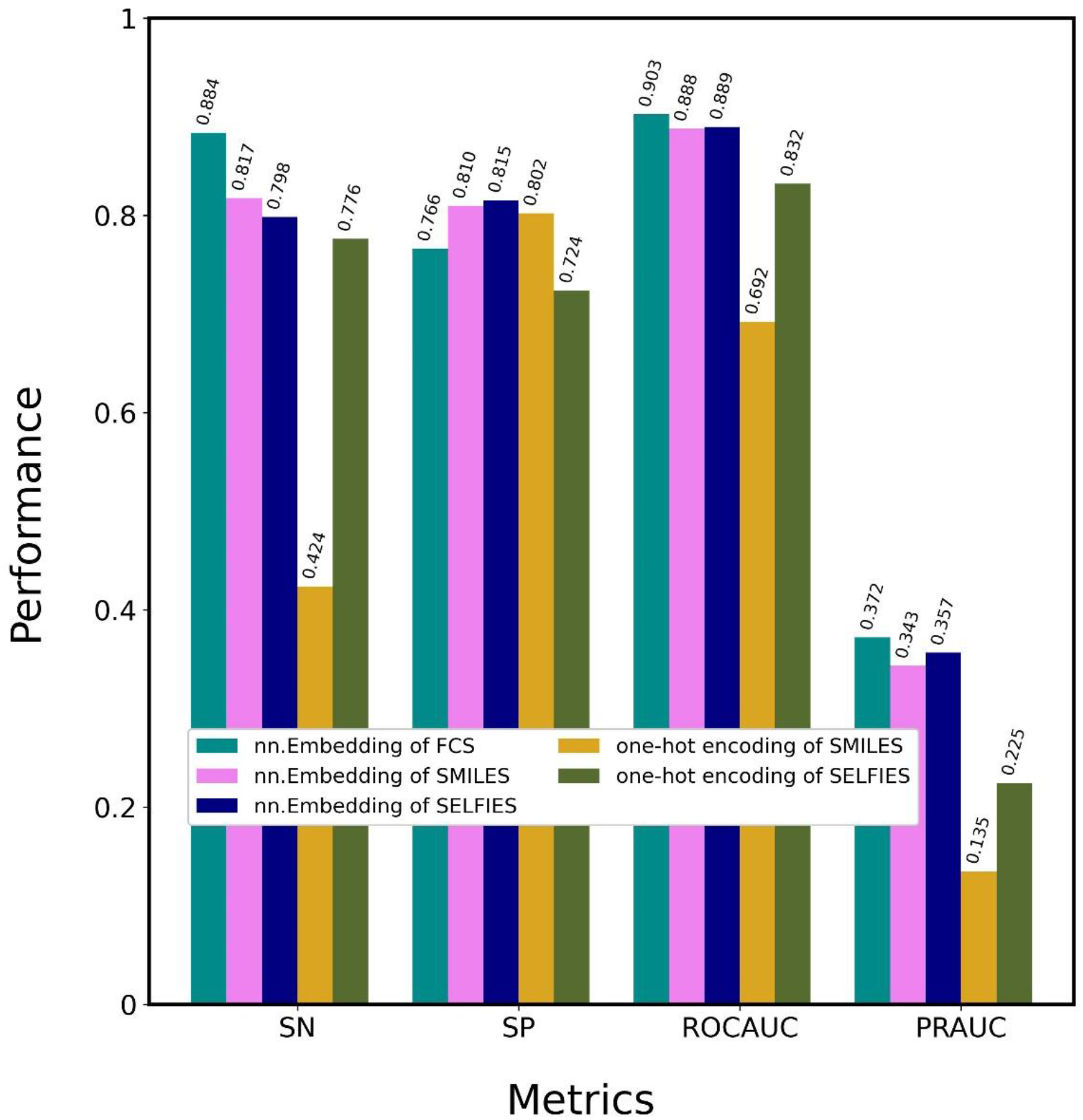
Optimization of encoding methods. CA_Ps (ICANs) with 5 different encoding methods were trained using the DAVIS training dataset and evaluated using the DAVIS test dataset. Details regarding the encoding methods are shown in Table 2.

### Comparison with state-of-the-art methods

We compared the optimal architecture of ICAN with 7 state-of-the-art methods on DAVIS, as shown in **Fig. 4** and **Table S3. Table S3** presents the values of all the six metrics including PR and F1. We employed a classical machine learning method, a linear regression (LR)-based model that used ECFP for encoding drugs and PSC for encoding proteins [32, 44], and the following CNN-based models: GNN-CPI [26], DeepDTI [9], DeepDTA [22], DeepConv-DTI [24], and the latest attention mechanism– based methods of TransformerCPI [13] and MolTrans [32]. ICAN and MolTrans outperformed the CNN-based methods (GNN-CPI, DeepDTI, DeepDTA, and DeepConv-DTI) in terms of ROCAUC and PRAUC. In particular, ICAN provided the highest SN, ROCAUC, and PRAUC values for all of the methods, although MolTrans was very competitive compared with ICAN. ICAN and MolTrans exhibited higher SN, SP, ROCAUC, and PRAUC values than TransformerCPI. The high PRAUC value of the ICAN method suggests that it takes advantage in the imbalanced dataset. The CNN-based methods provided higher SN, ROCAUC, and PRAUC values than the classical machine learning model of LR. In the CNN-based methods, DeepDTA exhibited the highest SN, whereas DeepConv-DTI exhibited the highest ROCAUC and PRAUC values.

**Figure 4.**
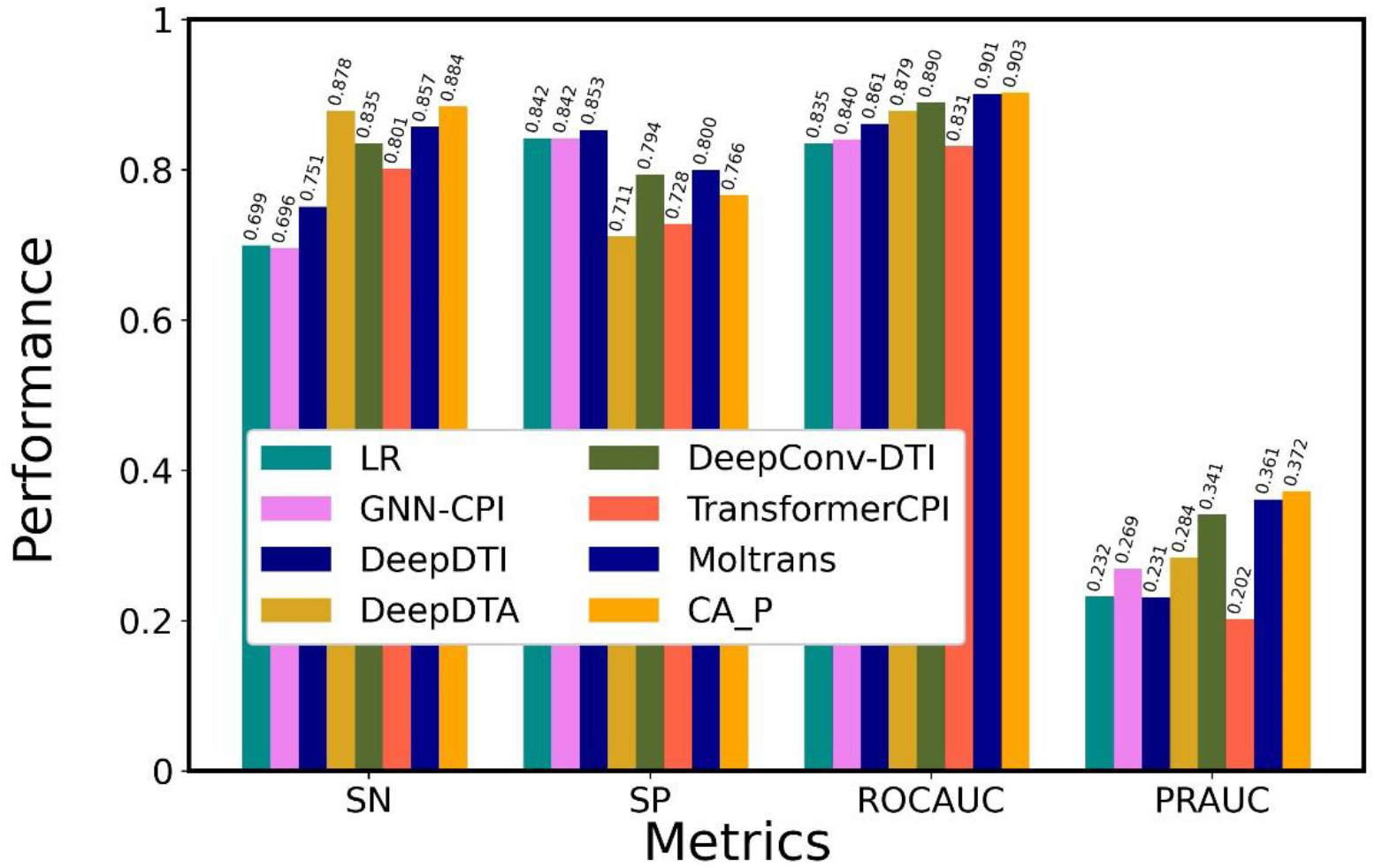
Comparison of the performance of ICAN with that of state-of-the-art models on DAVIS datasets. All models were trained using the DAVIS training dataset and evaluated using the DAVIS test dataset. CA_P was designated ICAN.

### Robustness

To characterize the robustness of the optimal network architecture of ICAN, we compared it with 7 state-of-the-art methods on the BindingDB and BIOSNAP datasets, as shown in **Fig.s 5** and **6** and **Tables S4** and **S5**. In BindingDB (**Fig. 5, Table S4**), ICAN provided the highest PRAUC value, whereas the CNN-based models (GNN-CPI, DeepDTA, and DeepConv-DTI) and MolTrans were competitive compared with ICAN in terms of ROCAUC and PRAUC values. For the three attention-based models (TransformerCPI, MolTrans, and ICAN), MolTrans and ICAN provided higher ROCAUC and PRAUC values than TransformerCPI. Although GNN-CPI provided high SP, ROCAUC, and PRAUC values, it exhibited poor performance on DAVIS. On BIOSNAP **(Fig. 6, Table S5**), the CNN-based methods (GNN-CPI, DeepDTI, DeepDTA, and DeepConv-DTI) provided high ROCAUC and PRAUC values that were competitive with or greater than those obtained with the three attention-based methods. DeepDTA provided the highest ROCAUC and PRAUC values for all of the methods. For the three attention-based models, MolTrans provided slightly higher ROCAUC and PRAUC values than ICAN, whereas TransformerCPI was competitive compared with ICAN. TransformerCPI functioned well using the balanced dataset (BIOSNAP), but it performed poorly on the imbalanced datasets (DAVIS, BindingDB).

**Figure 5.**
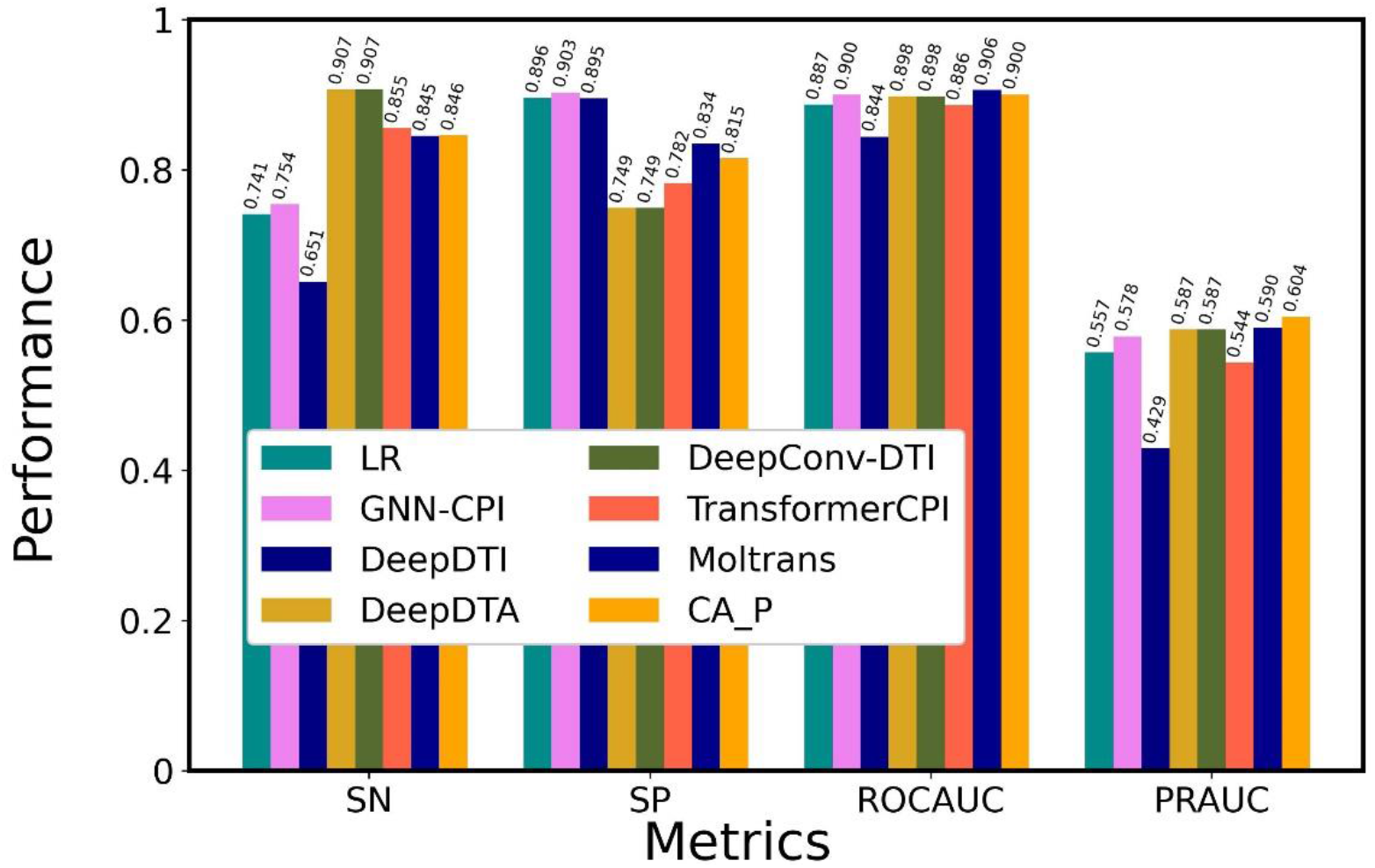
Comparison of the performance of ICAN with that of state-of-the-art models on BindingDB. All models were trained using the BindingDB training dataset and evaluated using the BindingDB test dataset. CA_P was designated ICAN.

**Figure 6.**
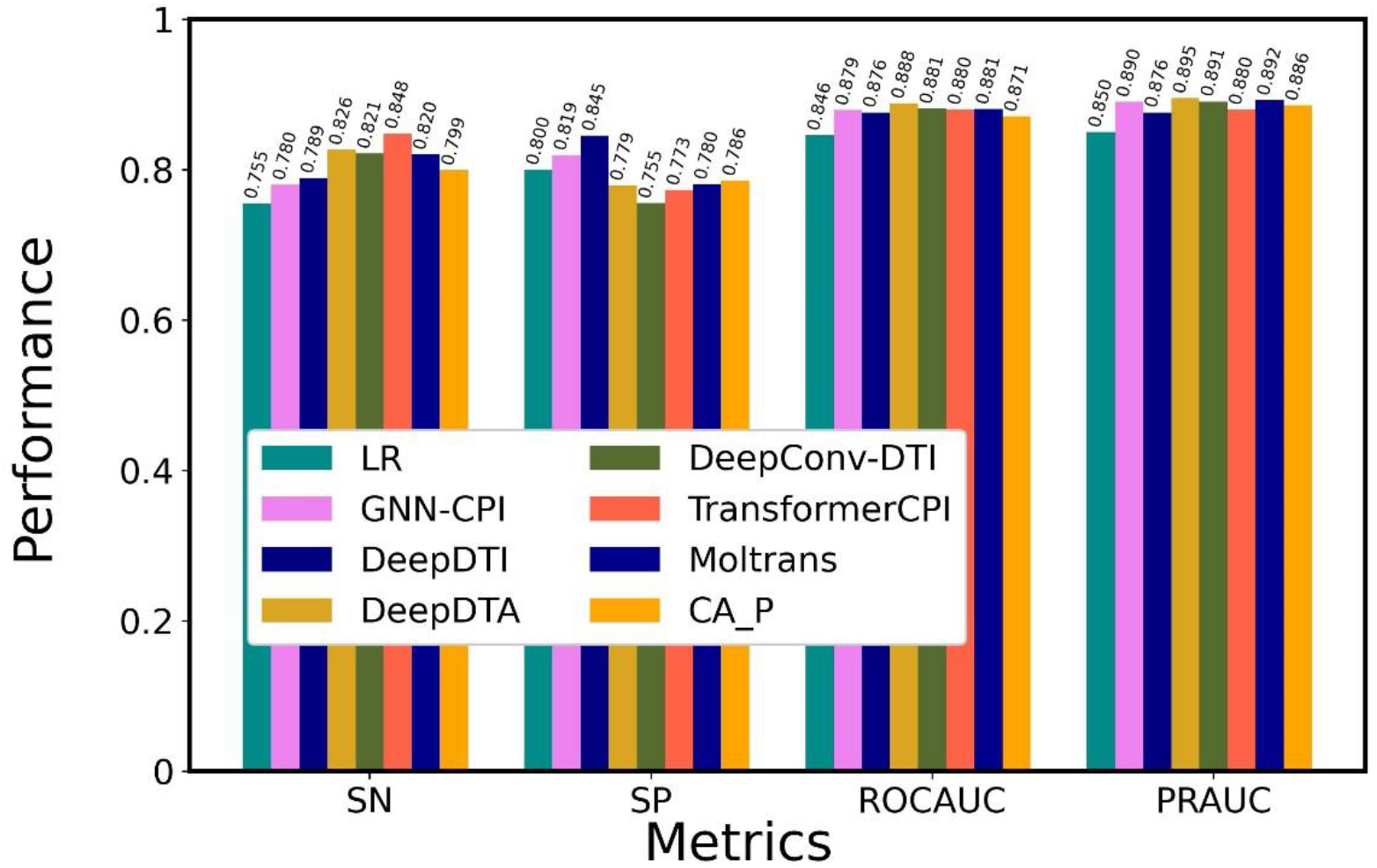
Comparison of the performance of ICAN with that of state-of-the-art models on BIOSNAP. All models were trained using the BIOSNAP training dataset and evaluated using the BIOSNAP test dataset. CA_P was designated ICAN.

The performance of the deep learning models including ICAN depended on the datasets. No method was ideal for all three datasets. ICAN provided the highest PRAUC value on the DAVIS and BindingDB test datasets (imbalanced dataset) but not on the BIOSNAP test dataset (balanced dataset). This suggests that ICNA has an advantage in predicting imbalanced test datasets. It is notable that the DAVIS and BindingDB test datasets contain a much greater abundance of negative samples than positive samples. In contrast, the BIOSNAP test dataset contains an equal number of negative and positive samples (**Table 1**). Both the attention-based methods (ICAN, MolTrans) and CNN-based methods (DeepDTA, DeepConv-DTI) exhibited competitive performance on the BindingDB test dataset. DeepDTA and DeepConv-DTI exhibited higher prediction performance on the BIOSNAP test dataset compared with the attention-based models. CNN may learn a large BIOSNAP dataset efficiently to provide high performance.

It is essential that DTI predictors exhibit generalization capability and robustness for external tests and practical applications. To this end, we must understand what deep learning models learn that can affect the generalization capability and use not only publically available data for drugs and proteins but also unlabeled biochemical and biophysical data, including structural information. In this regard, structural information emerging from the AlphaFold Protein Structure Database [45] would be useful for enabling accurate predictions.

### Feature visualization

We used principal component analysis to visualize how ICAN captures the features of positive and negative samples in the DAVIS test dataset, as shown in **Fig. 7**. Each sample consisted of concatenated feature matrixes of drugs and proteins. The matrixes embedded by nn.Embedding of FCS (**Fig. 7A**) were not separated between the positive and negative samples, with both sample types overlapping. The context matrixes resulting from the attention-mechanism (**Fig. 7B**) were somewhat separated compared with the concatenated embedding matrixes of the drugs and proteins. The attention mechanisms were shown to contribute to the classification of positive and negative samples, but additional improvements are needed for clear classification.

**Figure 7.**
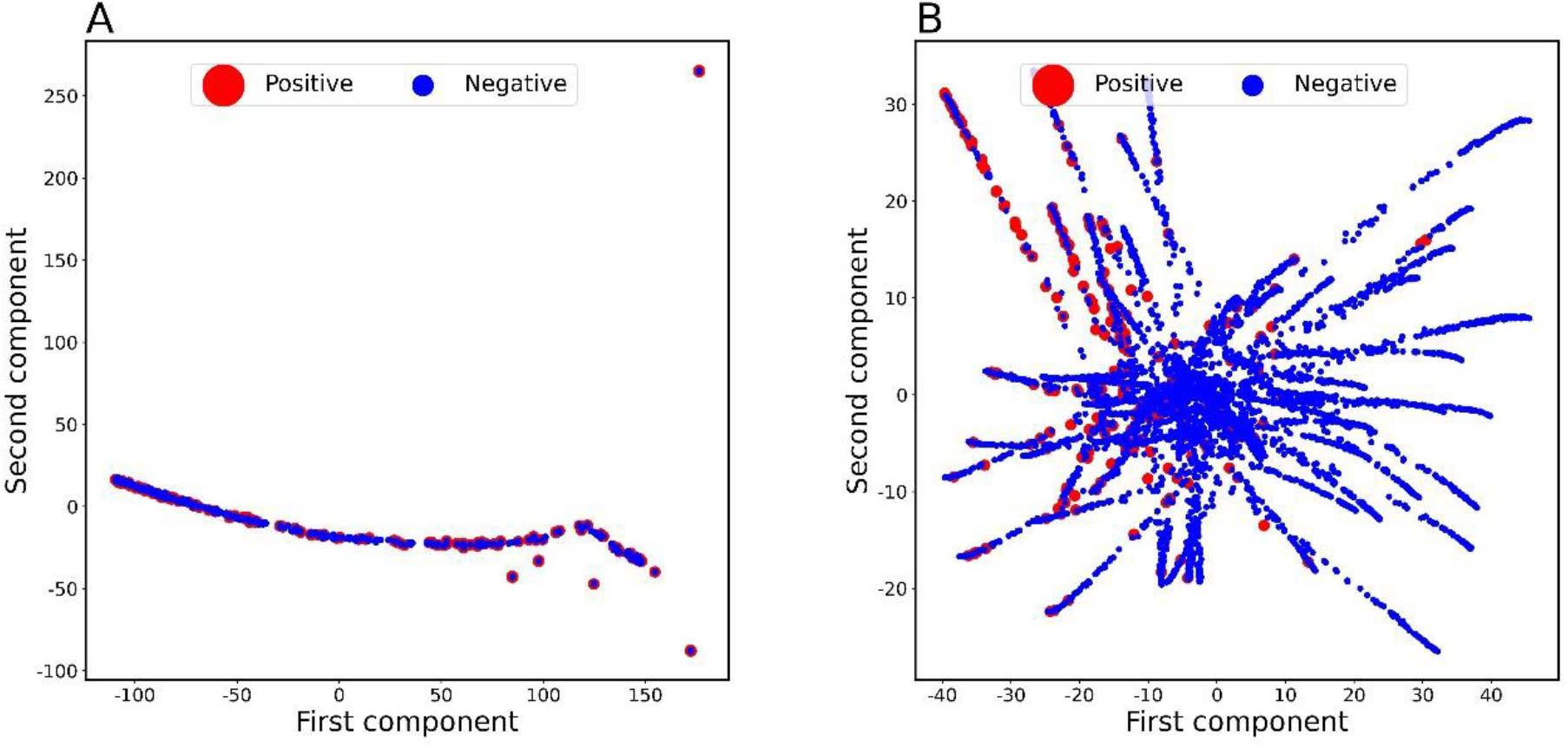
Visualization of feature matrixes for positive and negative samples on DAVIS. CA_P (ICAN) was tested on the DAVIS test dataset. (A) Feature matrixes embedded by nn.Embedding of FCS (B) Context matrixes generated by the attention mechanism.

### Interpretability of binding mechanism

To demonstrate the interpretability of the attention mechanisms, we fed the 160 DTIs with their binding sites into ICAN to generate corresponding attention weight matrixes (**Fig. 8A**) and calculated the experimental consistency number between the top 30 attention sites and the experimental drug-binding sites as 2.020 (**Fig. 8B**). We found that approximately 2 of the 30 attention sites corresponded to the experimental binding sites for each target protein. To statistically evaluate this number, we generated 10,000 simulated consistency numbers between the top 30 attention sites and randomly generated binding sites of proteins for 160 DTIs (**Fig. 8B**). The mean and standard deviation of the consistency number were 1.64 and 0.119, respectively. Based on this profile, the calculated z-score for the experimental consistency number (2.020) was 3.186, indicating that the attention sites were definitely related to the experimental binding sites.

**Figure 8.**
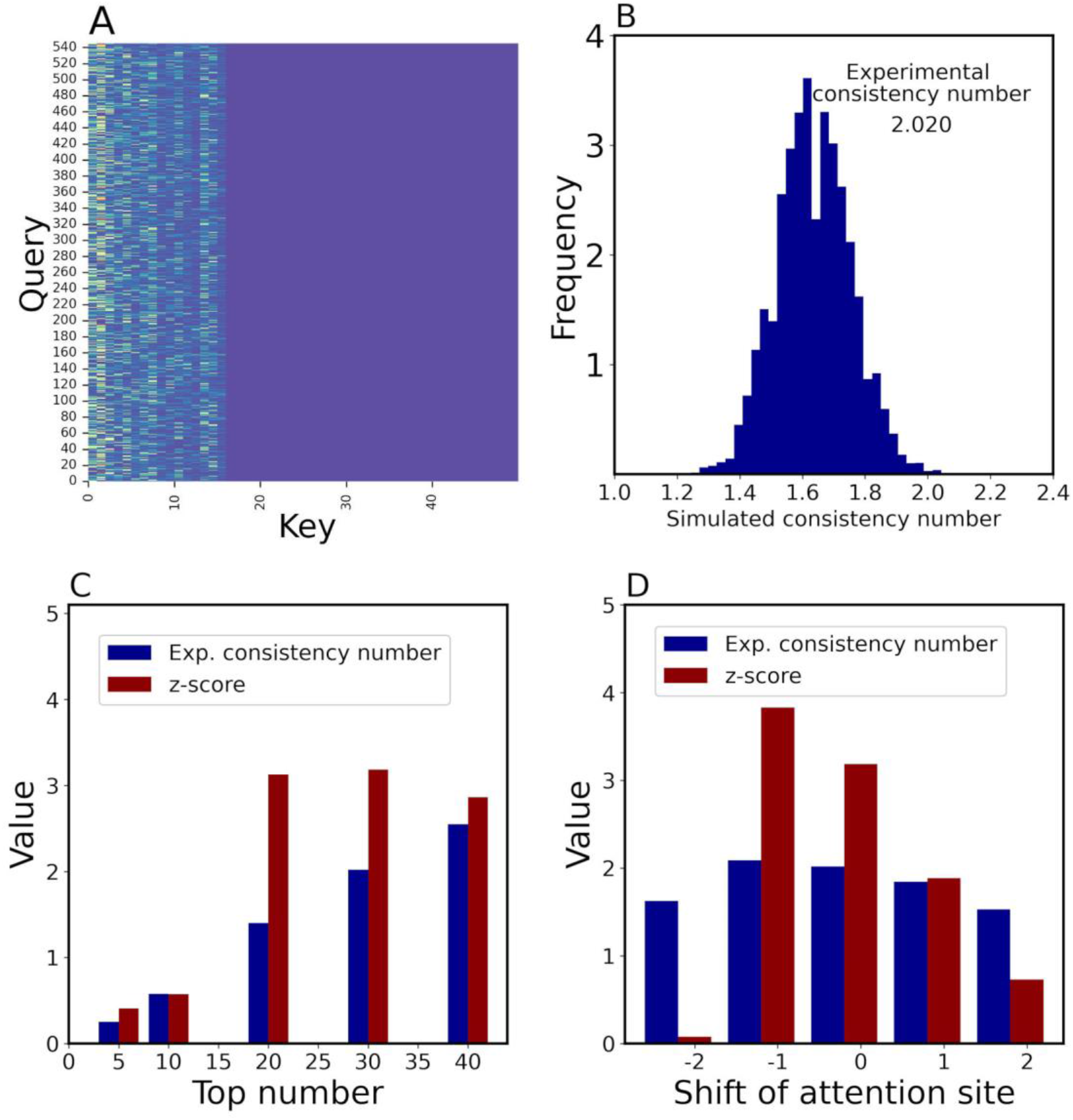
Attention-weight analysis. 160 DTIs with their binding sites were tested by ICAN. The simulation was iterated 10,000 times. (A) Attention-weight matrix for the DTI between DB06896 and P08581 (hepatocyte growth factor receptor). (B) Profile of simulated consistency numbers between the top 30 attention sites and randomly generated binding sites. (C) Effect of the number of top attention sites on experimental consistency numbers and associated z-scores. Experimental consistency numbers and associated z-scores were calculated with respect to a change in the top attention site number. (D) Effect of a shift of attention sites on experimental consistency number and their z-score values.

To determine why it is necessary to use the top 30 attention sites, we investigated how a change in the maximum number of attention sites affects the experimental consistency numbers and their corresponding z-scores, as shown in **Fig. 8C**. Using the top 30 attention sites provided the greatest increase in both the experimental consistency number and z-score. Thus, we selected the top 30 attention sites. Another reason was that the average number of experimental binding sites was approximately 28.9 (∼30) for each target protein. Using the top 10 attention sites provided a small experimental consistency number of 0.5, whereas the top 30 attention sites provided a number of 2. This suggests that the top-ranked attention site does not always represent the binding site. Only 2 of 30 attention sites corresponded to binding sites, whereas the remaining 28 did not. The remaining sites may interact indirectly with the binding sites or affect the biochemical and/or structural properties of DTIs. Consequently, although the binding sites are clearly focused by the attention mechanism, they are not priority factors of the mechanism. It would be very interesting to identify what the attention mechanism actually focuses on.

To further validate the relationship between attention sites and experimental binding sites, we shifted the attention sites by −2, −1, 0, 1, and 2 to calculate the experimental consistency numbers and corresponding z-scores. Zero indicates no shift of the attention sites, whereas a value of 1 indicates that the attention site was up-shifted by one amino acid. A shown in **Fig. 8C**, a shift of −1 did not reduce the experimental consistency number and its corresponding z-score, whereas shifts of −2, 1, and 2 decreased these values. These results demonstrate that the attention mechanism focuses on the exact binding site and neighboring sites.

Recently developed attention-based deep learning models have suggested that attention can be interpreted by investigating the relationship between highly weighted attention sites and experimental binding sites. Data from TransformerCPI suggested that weighted attention sites are related to binding sites of transforming growth factor-beta type 1 receptor and C-X-C chemokine receptor type 4 [13]. MolTrans suggested binding sites for Ephrin type-A receptor and Dasatinib and binding sites for histone deacetylase 2 and hydroxamic acid [32]. Although these studies provided a few successful examples, they did not conduct comprehensive statistical analyses of the results. To overcome this limitation, we first conducted a statistical analysis to determine the relationships between the attention sites and experimental binding sites and discovered that the attention sites were closely related to the experimental binding sites. This conclusion resulted from the plain structure of the cross-attention mechanism, which directly reads the drug and protein encoding features (**Fig. 1**).

## Conclusions

We designed an attention-based DTI method designated ICAN, which is composed of an embedding layer, attention mechanism, and output layer. Analyses of the attention mechanism factors (cross attention/self-attention, attention layer depth, selection of context matrixes from the attention networks, and output layer algorithms [CNN or fully connected layer]) revealed that the plain cross-attention mechanism with CNN and CA_P (ICAN) provided the highest prediction performance. The cross-attention mechanism considers sub-sequence interactions between a drug and a protein to produce context matrixes; the subsequent CNN extracts local sub-sequence patterns within the context matrixes using different filters. ICAN successfully decodes drug-related protein context features without the need for any protein-related drug context features. This is similar to TransformerCPI, which decodes protein features based on encoded drug features, but it is not consistent with MolTrans, which uses two self-attention mechanisms. MolTrans implements an interaction map that integrates the drug and protein context features from the self-attention mechanism to complement the lack of a cross-attention mechanism between drugs and proteins. We then selected nn.Embedding of FCS as the best of the five encoding methods: nn.Embedding of FCS, nn.Embedding of SMILES, nn.Embedding of SELFIES, one-hot encoding of SMILES, and one-hot encoding of SELFIES. Furthermore, we found that SELFIES was more effective than SMILES in terms of DTI prediction, because SELFIES decomposes chemical formulas into a chemically meaningful character list, but SMILES does not.

We also characterized the optimal architecture of ICAN in comparison with 7 state-of-the-art methods: a classical LR-based method, CNN-based models (GNN-CPI, DeepDTI, DeepDTA, and DeepConv-DTI), and the latest attention mechanism–based methods (TransformerCPI, MolTrans). ICAN provided the highest PRAUC value on the imbalanced datasets (DAVIS, BindingDB) but not on the balanced dataset (BIOSNAP). These results suggest that ICAN has an advantage in predicting imbalanced test datasets. The attention-based methods provided higher ROCAUC and PRAUC values than CNN-based models on DAVIS. Both the attention-based methods (ICAN and MolTrans) and CNN-based methods (DeepDTA and DeepConv-DTI) showed very competitive performance on BindingDB. DeepDTA and DeepConv-DTI exhibited better performance than the attention-based models on BIOSNAP. No best method was identified for all three datasets. These results indicate that the performance of deep learning models depends on the dataset.

The most important task is to first conduct a comprehensive statistical analysis to clearly determine the relationships between attention sites and experimental binding sites. ICAN was shown to enable the interpretation of various DTI mechanisms. We consider that such interpretability results from the plain structure of the cross-attention mechanism between a drug and protein. On the other hand, our analyses also suggested that the attention mechanism captures critical factors other than binding sites, as it was shown to explain only a few of the 30 attention sites considered. To further increase the interpretability and robustness in examining independent datasets, it is necessary to determine exactly what the attention mechanism really captures and how this affects the generalization capability. Additional DTI predictors remain to be developed using different chemo-genomics approaches.

## Data availability

The program is freely available at https://github.com/kuratahiroyuki/ICAN.git.

## Acknowledgments

We are grateful to Mitsuki Watanabe for collecting DTI binding-site data. This work was supported by a Grant-in-Aid for Scientific Research (B) (22H03688) and partially supported by a Grant-in-Aid for JSPS Research Fellows (22J22706) from the Japan Society for the Promotion of Science (JSPS).

## References

1. Broach JR, Thorner J. High-throughput screening for drug discovery, Nature 1996;384:14–16.

2. Ezzat A, Wu M, Li XL et al. Computational prediction of drug-target interactions using chemogenomic approaches: an empirical survey, Brief Bioinform 2019;20:1337–1357.

3. Yamanishi Y, Araki M, Gutteridge A et al. Prediction of drug-target interaction networks from the integration of chemical and genomic spaces, Bioinformatics 2008;24:i232–240.

4. Ding H, Takigawa I, Mamitsuka H et al. Similarity-based machine learning methods for predicting drug-target interactions: a brief review, Brief Bioinform 2014;15:734–747.

5. Pahikkala T, Airola A, Pietila S et al. Toward more realistic drug-target interaction predictions, Brief Bioinform 2015;16:325–337.

6. Gonen M. Predicting drug-target interactions from chemical and genomic kernels using Bayesian matrix factorization, Bioinformatics 2012;28:2304–2310.

7. Zheng X. Collaborative matrix factorization with multiple similarities for predicting drug-target interactions. In: KDD. Chicago, 2013.

8. Ezzat A, Zhao P, Wu M et al. Drug-Target Interaction Prediction with Graph Regularized Matrix Factorization, IEEE/ACM Trans Comput Biol Bioinform 2017;14:646–656.

9. Wen M, Zhang Z, Niu S et al. Deep-Learning-Based Drug-Target Interaction Prediction, J Proteome Res 2017;16:1401–1409.

10. Rogers D, Hahn M. Extended-connectivity fingerprints, J Chem Inf Model 2010;50:742–754.

11. Bolton EE. Pubchem: integrated platform of small molecules and biological activities. Annual Reports in Computational Chemistry. Elsevier, 2008, 217–241.

12. Dubchak I, Muchnik I, Holbrook SR et al. Prediction of protein folding class using global description of amino acid sequence, Proc Natl Acad Sci U S A 1995;92:8700–8704.

13. Chen L, Tan X, Wang D et al. TransformerCPI: improving compound-protein interaction prediction by sequence-based deep learning with self-attention mechanism and label reversal experiments, Bioinformatics 2020;36:4406–4414.

14. Yu H, Chen J, Xu X et al. A systematic prediction of multiple drug-target interactions from chemical, genomic, and pharmacological data, PLoS One 2012;7:e37608.

15. Bagherian M, Sabeti E, Wang K et al. Machine learning approaches and databases for prediction of drug-target interaction: a survey paper, Brief Bioinform 2021;22:247–269.

16. He T, Heidemeyer M, Ban F et al. SimBoost: a read-across approach for predicting drug-target binding affinities using gradient boosting machines, J Cheminform 2017;9:24.

17. Ezzat A, Wu M, Li XL et al. Drug-target interaction prediction via class imbalance-aware ensemble learning, BMC Bioinformatics 2016;17:509.

18. Islam SM, Hossain SMM, Ray S. DTI-SNNFRA: Drug-target interaction prediction by shared nearest neighbors and fuzzy-rough approximation, PLoS One 2021;16:e0246920.

19. Hamanaka M, Taneishi K, Iwata H et al. CGBVS-DNN: Prediction of Compound-protein Interactions Based on Deep Learning, Mol Inform 2017;36.

20. Yu L, Qiu W, Lin W et al. HGDTI: predicting drug-target interaction by using information aggregation based on heterogeneous graph neural network, BMC Bioinformatics 2022;23:126.

21. Abbasi K, Razzaghi P, Poso A et al. DeepCDA: deep cross-domain compound-protein affinity prediction through LSTM and convolutional neural networks, Bioinformatics 2020;36:4633–4642.

22. Ozturk H, Ozgur A, Ozkirimli E. DeepDTA: deep drug-target binding affinity prediction, Bioinformatics 2018;34:i821–i829.

23. Sutskever I, Hinton GE. Deep, narrow sigmoid belief networks are universal approximators, Neural Comput 2008;20:2629–2636.

24. Lee I, Keum J, Nam H. DeepConv-DTI: Prediction of drug-target interactions via deep learning with convolution on protein sequences, PLoS Comput Biol 2019;15:e1007129.

25. Karimi M, Wu D, Wang Z et al. DeepAffinity: interpretable deep learning of compound-protein affinity through unified recurrent and convolutional neural networks, Bioinformatics 2019;35:3329–3338.

26. Tsubaki M, Tomii K, Sese J. Compound-protein interaction prediction with end-to-end learning of neural networks for graphs and sequences, Bioinformatics 2019;35:309–318.

27. Nguyen T, Le H, Quinn TP et al. GraphDTA: predicting drug-target binding affinity with graph neural networks, Bioinformatics 2021;37:1140–1147.

28. Scarselli F, Gori M, Tsoi AC et al. The graph neural network model, IEEE Trans Neural Netw 2009;20:61–80.

29. Lim J, Ryu S, Park K et al. Predicting Drug-Target Interaction Using a Novel Graph Neural Network with 3D Structure-Embedded Graph Representation, J Chem Inf Model 2019;59:3981–3988.

30. Kipf T, Welling M. Semi-supervised classification with graph convolutional networks. 5th International Conference on Learning Representations. 2017, 1–14.

31. Torng W, Altman RB. Graph Convolutional Neural Networks for Predicting Drug-Target Interactions, J Chem Inf Model 2019;59:4131–4149.

32. Huang K, Xiao C, Glass LM et al. MolTrans: Molecular Interaction Transformer for drug-target interaction prediction, Bioinformatics 2021;37:830–836.

33. Davis MI, Hunt JP, Herrgard S et al. Comprehensive analysis of kinase inhibitor selectivity, Nat Biotechnol 2011;29:1046–1051.

34. Liu T, Lin Y, Wen X et al. BindingDB: a web-accessible database of experimentally determined protein-ligand binding affinities, Nucleic Acids Res 2007;35:D198–201.

35. Zitnik M, Sosi R, Maheshwari S et al. BioSNAP datasets: Stanford biomedical network dataset collection., https://snap.stanford.edu/biodata/index.html 2018.

36. Wishart DS, Knox C, Guo AC et al. DrugBank: a knowledgebase for drugs, drug actions and drug targets, Nucleic Acids Res 2008;36:D901–906.

37. Landrum G. RDKit: Open-source cheminformatics, https://www.rdkit.org 2006;3.

38. Weininger D. SMILES, a chemical language and information system. 1. Introduction to methodology and encoding rules, J. Chem. Inf. Comput. Sci. 1988;28:31–36.

39. Krenn M, Haese F, AkshatKumar Nigam et al. Self-Referencing Embedded Strings (SELFIES): A 100% robust molecular string representation, Machine Learning: Science and Technology 2020;1:045024.

40. Sennrich R, Haddow B, Birch A. Neural Machine Translation of Rare Words with Subword Units. 2016, p. 1715–1725. Association for Computational Linguistics.

41. Gage P. A new algorithm for data compression, C Users J. 1994;12:23–38.

42. Meslamani J, Rognan D, Kellenberger E. sc-PDB: a database for identifying variations and multiplicity of ‘druggable’ binding sites in proteins, Bioinformatics 2011;27:1324–1326.

43. Paszke A, Gross S, Massa F et al. PyTorch: An Imperative Style, High-Performance Deep Learning Library, 33rd Conference on Neural Information Processing Systems (NeurIPS 2019), Vancouver, Canada 2019:1–12.

44. Cao DS, Xu QS, Liang YZ. propy: a tool to generate various modes of Chou’s PseAAC, Bioinformatics 2013;29:960–962.

45. Jumper J, Evans R, Pritzel A et al. Highly accurate protein structure prediction with AlphaFold, Nature 2021;596:583–589.

