## Supplemental information for "ICAN: interpretable cross-attention network for identifying drug and target protein interactions"

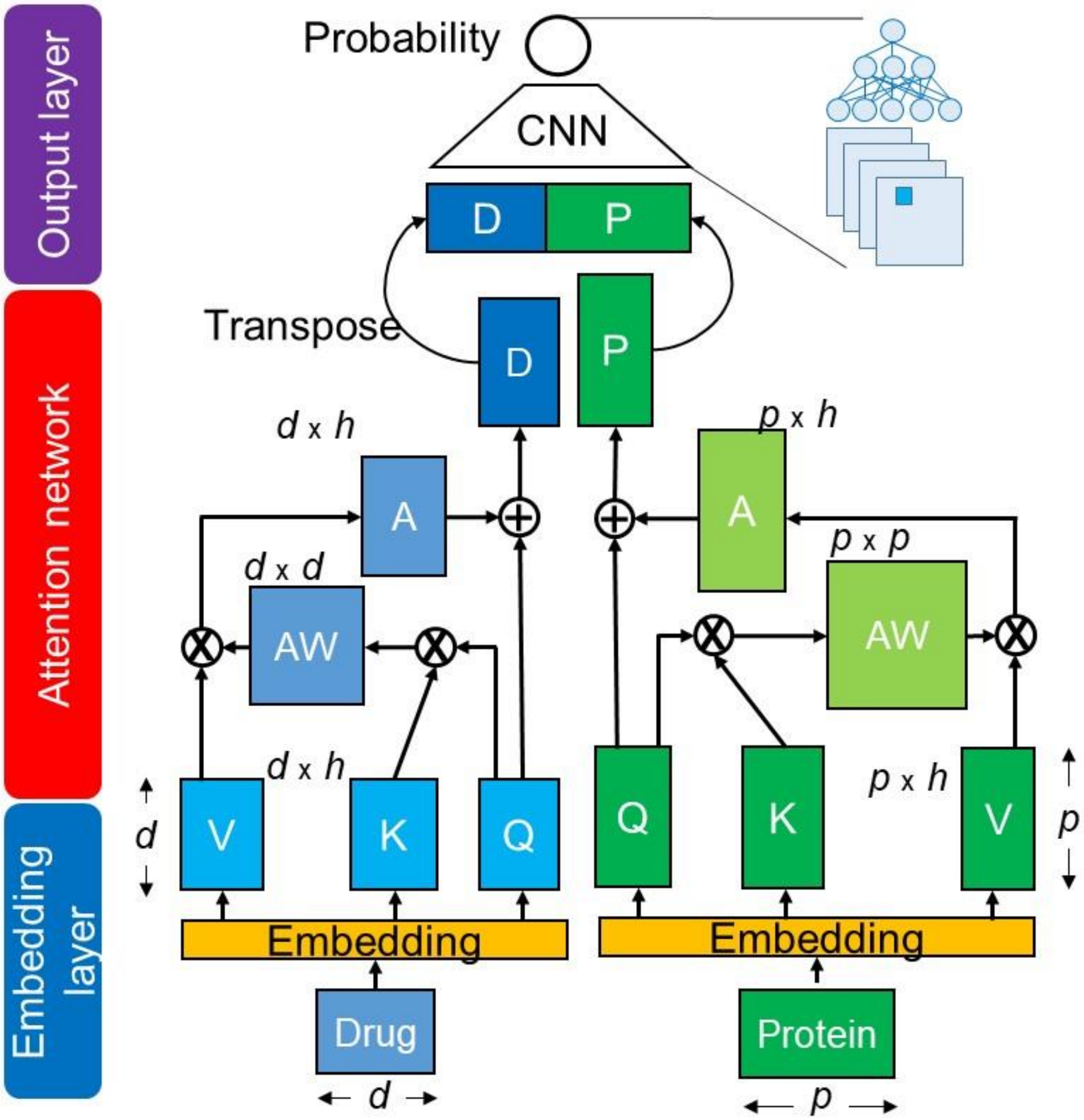


**Figure S1 Self-attention–based model**

This network corresponds to SA_DP with nn.Embedding of FCS (Table 3). Q, K, and V denote Query, Key, and Value matrixes.

AW: attention-weight matrix, A: attention matrix, D: drug-context matrix, P: protein-context matrix, d: length of drug sequence, p: length of protein sequence, h: hidden dimension size.

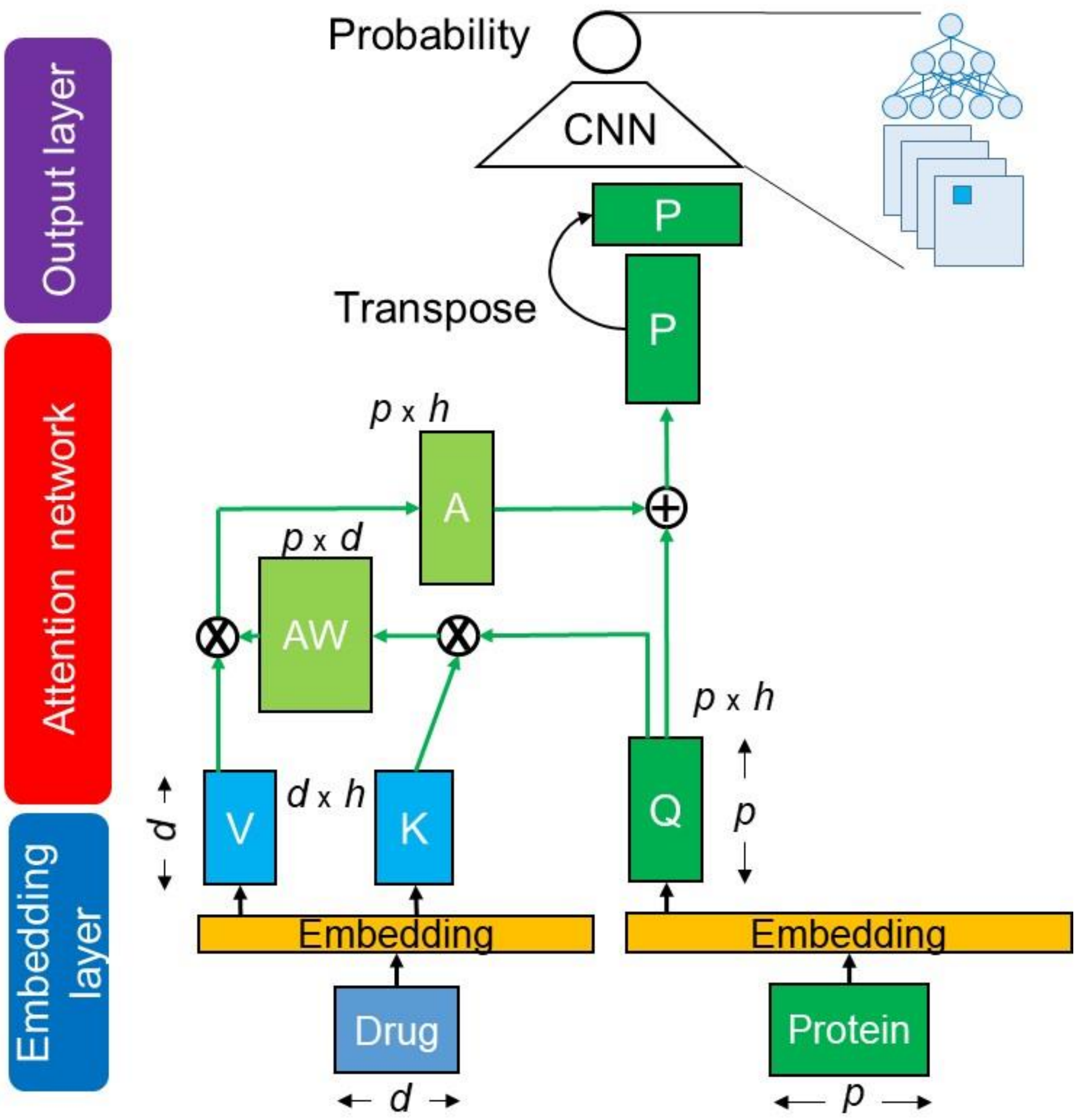


**Figure S2 Cross-attention–based model**

This network corresponds to CA_P with nn.Embedding of FCS (Table 3). It uses only protein context features. Q, K, and V denote Query, Key, and Value matrixes.

AW: attention-weight matrix, A: attention matrix, D: drug-context matrix, P: protein-context matrix, d: length of drug sequence, p: length of protein sequence, h: hidden dimension size.

**Table S1 Performance of different network architectures**

| Architecture | | SN | SP | ROCAUC | PR | F1 | PRAUC |
|---|---|---|---|---|---|---|---|
| CA_DP_FCL | Mean | 0.802 | **0.814** | 0.890 | **0.186** | **0.302** | 0.359 |
| | Std | 0.012 | 0.009 | 0.004 | 0.007 | 0.009 | 0.017 |
| CA_DP | Mean | 0.843 | 0.789 | 0.897 | 0.183 | 0.299 | 0.367 |
| | Std | 0.044 | 0.066 | 0.006 | 0.038 | 0.048 | 0.016 |
| CA2_DP | Mean | 0.826 | 0.793 | 0.884 | 0.179 | 0.293 | 0.314 |
| | Std | 0.044 | 0.043 | 0.010 | 0.025 | 0.031 | 0.011 |
| CA3_DP | Mean | 0.833 | 0.777 | 0.880 | 0.166 | 0.277 | 0.284 |
| | Std | 0.029 | 0.019 | 0.007 | 0.009 | 0.012 | 0.036 |
| SA_DP | Mean | 0.818 | 0.710 | 0.840 | 0.131 | 0.226 | 0.243 |
| | Std | 0.034 | 0.040 | 0.008 | 0.012 | 0.017 | 0.013 |
| CA_D | Mean | 0.799 | 0.812 | 0.879 | 0.186 | 0.301 | 0.306 |
| | Std | 0.023 | 0.026 | 0.008 | 0.017 | 0.021 | 0.014 |
| CA_P (ICAN) | Mean | **0.884** | 0.766 | **0.903** | 0.167 | 0.281 | **0.372** |
| | Std | 0.011 | 0.016 | 0.005 | 0.009 | 0.012 | 0.032 |

PR denotes precision. F1 denotes F1-score that is the harmonic mean of PR and recall (SP). Mean and Std denote the mean and standard deviation of each metric. The architectures are shown in Table 3. Bold values indicate the best-performing method for each metric.

**Table S2 Performance of different encoding methods in CA_P (ICAN)**

| Encoding method | | SN | SP | ROCAUC | PR | F1 | PRAUC |
|---|---|---|---|---|---|---|---|
| nn.Embedding of FCS | Mean | **0.884** | 0.766 | **0.903** | 0.167 | 0.281 | **0.372** |
| | Std | 0.011 | 0.016 | 0.005 | 0.009 | 0.012 | 0.032 |
| nn.Embedding of SMILES | Mean | 0.817 | 0.810 | 0.888 | **0.189** | **0.306** | 0.343 |
| | Std | 0.044 | 0.039 | 0.014 | 0.026 | 0.033 | 0.033 |
| nn.Embedding of SELFIES | Mean | 0.798 | **0.815** | 0.889 | 0.188 | 0.304 | 0.357 |
| | Std | 0.045 | 0.029 | 0.008 | 0.014 | 0.016 | 0.026 |
| One-hot encoding of SMILES | Mean | 0.424 | 0.802 | 0.692 | 0.063 | 0.109 | 0.135 |
| | Std | 0.404 | 0.194 | 0.101 | 0.060 | 0.104 | 0.064 |
| One-hot encoding of SELFIES | Mean | 0.776 | 0.724 | 0.832 | 0.151 | 0.246 | 0.225 |
| | Std | 0.080 | 0.142 | 0.034 | 0.059 | 0.078 | 0.041 |

PR denotes precision. F1 denotes F1-score that is the harmonic mean of PR and recall (SP). Mean and Std denote the mean and standard deviation of each metric. The encoding methods are shown in Table 2. Bold values indicate the best-performing method for each metric.

**Table S3 Performance of different learning methods on the DAVIS test dataset**

| Method | | SN | SP | ROCAUC | PR | F1 | PRAUC |
|---|---|---|---|---|---|---|---|
| LR | Mean | 0.699 | 0.842 | 0.835 | - | - | 0.232 |
| | Std | 0.051 | 0.033 | 0.010 | - | - | 0.023 |
| GNN-CPI | Mean | 0.696 | 0.842 | 0.840 | - | - | 0.269 |
| | Std | 0.047 | 0.039 | 0.012 | - | - | 0.020 |
| DeepDTI | Mean | 0.751 | **0.853** | 0.861 | - | - | 0.231 |
| | Std | 0.015 | 0.012 | 0.002 | - | - | 0.006 |
| DeepDTA | Mean | 0.878 | 0.711 | 0.879 | 0.140 | 0.242 | 0.284 |
| | Std | 0.023 | 0.040 | 0.008 | 0.013 | 0.019 | 0.022 |
| DeepConv-DTI | Mean | 0.835 | 0.794 | 0.890 | 0.180 | 0.295 | 0.341 |
| | Std | 0.036 | 0.039 | 0.014 | 0.023 | 0.032 | 0.041 |
| TransformerCPI | Mean | 0.801 | 0.728 | 0.831 | 0.135 | 0.231 | 0.202 |
| | Std | 0.023 | 0.020 | 0.006 | 0.006 | 0.009 | 0.007 |
| MolTrans | Mean | 0.857 | 0.800 | 0.901 | **0.185** | **0.304** | 0.361 |
| | Std | 0.003 | 0.001 | 0.001 | 0.002 | 0.002 | 0.003 |
| CA_P (ICAN) | Mean | **0.884** | 0.766 | **0.903** | 0.167 | 0.281 | **0.372** |
| | Std | 0.011 | 0.016 | 0.005 | 0.009 | 0.012 | 0.032 |

PR denotes precision. F1 denotes F1-score that is the harmonic mean of PR and recall (SP). Mean and Std denote the mean and standard deviation of each metric. Bold values indicate the best-performing method for each metric.

**Table S4 Performance of different learning methods on the BindingDB test dataset**

| Method | | SN | SP | ROCAUC | PR | F1 | PRAUC |
|---|---|---|---|---|---|---|---|
| LR | Mean | 0.741 | 0.896 | 0.887 | - | - | 0.557 |
| | Std | 0.013 | 0.011 | 0.002 | - | - | 0.015 |
| GNN-CPI | Mean | 0.754 | **0.903** | 0.900 | - | - | 0.578 |
| | Std | 0.015 | 0.011 | 0.004 | - | - | 0.015 |
| DeepDTI | Mean | 0.651 | 0.895 | 0.844 | - | - | 0.429 |
| | Std | 0.024 | 0.023 | 0.002 | - | - | 0.005 |
| DeepDTA | Mean | **0.907** | 0.749 | 0.898 | 0.385 | 0.537 | 0.587 |
| | Std | 0.043 | 0.070 | 0.034 | 0.054 | 0.047 | 0.132 |
| DeepConv-DTI | Mean | **0.907** | 0.749 | 0.898 | 0.385 | 0.537 | 0.587 |
| | Std | 0.043 | 0.070 | 0.034 | 0.054 | 0.047 | 0.132 |
| TransformerCPI | Mean | 0.855 | 0.782 | 0.886 | 0.398 | 0.542 | 0.544 |
| | Std | 0.021 | 0.028 | 0.002 | 0.022 | 0.018 | 0.008 |
| MolTrans | Mean | 0.845 | 0.834 | **0.906** | **0.462** | **0.597** | 0.590 |
| | Std | 0.006 | 0.014 | 0.003 | 0.019 | 0.015 | 0.005 |
| CA_P (ICAN) | Mean | 0.846 | 0.815 | 0.900 | 0.434 | 0.574 | **0.604** |
| | Std | 0.023 | 0.015 | 0.003 | 0.014 | 0.008 | 0.016 |

PR denotes precision. F1 denotes F1-score that is the harmonic mean of PR and recall (SP). Mean and Std denote the mean and standard deviation of each metric. Bold values indicate the best-performing method for each metric.

**Table S5 Performance of different learning methods on the BIOSNAP test dataset**

| Method | | SN | SP | ROCAUC | PR | F1 | PRAUC |
|---|---|---|---|---|---|---|---|
| LR | Mean | 0.755 | 0.800 | 0.846 | - | - | 0.850 |
| | Std | 0.039 | 0.018 | 0.004 | - | - | 0.011 |
| GNN-CPI | Mean | 0.780 | 0.819 | 0.879 | - | - | 0.890 |
| | Std | 0.014 | 0.012 | 0.007 | - | - | 0.004 |
| DeepDTI | Mean | 0.789 | **0.845** | 0.876 | - | - | 0.876 |
| | Std | 0.027 | 0.017 | 0.005 | - | - | 0.006 |
| DeepDTA | Mean | 0.826 | 0.779 | **0.888** | **0.797** | 0.809 | **0.895** |
| | Std | 0.051 | 0.088 | 0.006 | 0.051 | 0.012 | 0.005 |
| DeepConv-DTI | Mean | 0.821 | 0.755 | 0.881 | 0.780 | 0.796 | 0.891 |
| | Std | 0.081 | 0.095 | 0.006 | 0.055 | 0.018 | 0.007 |
| TransformerCPI | Mean | **0.848** | 0.773 | 0.880 | 0.791 | **0.818** | 0.880 |
| | Std | 0.003 | 0.008 | 0.002 | 0.006 | 0.003 | 0.003 |
| MolTrans | Mean | 0.820 | 0.780 | 0.881 | 0.791 | 0.805 | 0.892 |
| | Std | 0.001 | 0.000 | 0.000 | 0.000 | 0.000 | 0.000 |
| CA_P (ICAN) | Mean | 0.799 | 0.786 | 0.871 | 0.791 | 0.795 | 0.886 |
| | Std | 0.012 | 0.018 | 0.001 | 0.012 | 0.003 | 0.002 |

PR denotes precision. F1 denotes F1-score that is the harmonic mean of PR and recall (SP). Mean and Std denote the mean and standard deviation of each metric. Bold values indicate the best-performing method for each metric.
